# High Efficiency Recombinant Protein Purification Using mCherry and YFP Nanobody Affinity Matrices

**DOI:** 10.1101/2021.12.01.470687

**Authors:** Anh T.Q. Cong, Taylor L. Witter, Matthew J. Schellenberg

**Affiliations:** Department of Biochemistry and Molecular Biology, Mayo Clinic, Rochester, MN, USA 55905

**Keywords:** recombinant protein expression, protein purification, single-domain antibody (sdAb, nanobody), protein structure, protein-protein interaction

## Abstract

Mammalian cell lines are important expression systems for large proteins and protein complexes, particularly when the acquisition of post-translational modifications in the protein’s native environment is desired. However, low or variable transfection efficiencies are challenges that must be overcome to use such an expression system. Expression of recombinant proteins as a fluorescent protein fusion enables real-time monitoring of protein expression, and also provides an affinity handle for one-step protein purification using a suitable affinity reagent. Here we describe a panel of anti-GFP and anti-mCherry nanobody affinity matrices and their efficacy for purification of GFP/YFP or mCherry fusion proteins. We define the molecular basis by which they bind their target protein using X-ray crystallography. From these analyses we define an optimal pair of nanobodies for purification of recombinant protein tagged with GFP/YFP or mCherry, and demonstrate these nanobody-sepharose supports are stable to many rounds of cleaning and extended incubation in denaturing conditions. Finally, we demonstrate the utility of the mCherry-tag system by using it to purify recombinant human Topoisomerase 2α expressed in HEK293F cells. The mCherry-tag and GFP/YFP-tag expression systems can be utilized for recombinant protein expression individually or in tandem for mammalian protein expression systems where real-time monitoring of protein expression levels and a high-efficiency purification step is needed.

## Introduction

Recombinant proteins are a central pillar of research in life sciences and drug discovery and are additionally beneficial as therapeutics to treat disease (1). Proteins are polypeptide chains that must achieve the correct 3–dimensional folding state. While some proteins may contain intrinsically disordered regions that are important for their function (2), globular proteins that fail to fold into their correct 3-dimensional structure are non-functional. When generating recombinant human proteins, bacterial protein expression systems are typically a first choice, but it can be challenging to obtain correctly–folded protein using this heterologous system (3). An alternative expression system is eukaryotic cells that can be grown in liter-scale suspension cultures. HEK293F is a human cell line that grows in suspension culture and can be transfected with recombinant DNA that encodes the protein of interest. As a human cell line, HEK293F cells also contain a cellular environment with the correct chaperones, post-translational modification machinery, and protein quality control systems for human recombinant protein expression. However, they have a more complex proteome than *Escherichia coli* which can complicate purification, and the use of transfection–based transgene delivery necessitates careful optimization of transfection conditions (4).

Methods for rapidly measuring protein expression and purifying recombinant protein from cell lysates are needed to effectively utilize mammalian expression systems. Recombinant proteins are often expressed fused to an affinity tag that can be used for initial purification from cell lysate. A hexa-histidine (His_6_) affinity tag with Ni-NTA resin is used widely for purification of proteins expressed in *E. coli*. However, His_6_ binding to a Ni-NTA affinity support can be weaker (17–700 nM K_d_) than other affinity matrices and has unfavorable dissociation kinetics that can result in sub-optimal yields of proteins that express at low levels (5,6). Furthermore, proteins can bind non-specifically to Ni-NTA, as some endogenous mammalian proteins contain His_≥6_ sequences necessitating the use of affinity tags and solid-phase supports with higher affinity and specificity. Other tags such as maltose binding protein (MBP) has a similar binding affinity (660 nM K_d_) (7) or glutathione S-transferase (8), which binds more tightly to its ligand (7 nM K_d_) cannot be directly visualized.

The Yellow Fluorescent Protein (YFP)–tag expression system overcomes two key challenges to HEK293F–based recombinant protein expression (9). The first is that YFP fluorescence can be monitored to provide a real-time measurement of protein expression levels, so transfection conditions, post-transfection growth time, and binding to an affinity resin can be monitored and optimized without time-consuming steps such as western blotting. The second is that YFP can also be used as an affinity tag for a single high-efficiency purification step using an anti-Green Fluorescent Protein (GFP) nanobody (which also binds the nearly identical YFP) coupled to sepharose resin. Although anti-GFP/YFP antibodies can be used for recombinant protein purification (10), it is much more feasible to use nanobodies, which are the ∼14 kDa VHH domain fragment of camelid single-chain antibodies that possess the entire antigen binding site and typically have nanomolar affinity for their antigen (11). Importantly, the GFP-nanobody has very slow dissociation kinetics (1.45 × 10^-4^ s^-1^ (12)) so it can retain YFP-tagged protein for the extended durations needed to recover low concentrations of recombinant protein from cell lysates. Nanobodies against fluorescent proteins (FP)s have been chemically linked to sepharose resin to generate an affinity support for immunoprecipitation of FP-tagged proteins (12-14). In this way the GFP-enhancer nanobody has been used for purification of YFP-tagged recombinant Topoisomerase 2 and ZATT proteins (14,15). The effectiveness of the YFP-tag approach in producing these proteins warrants further investigation to identify the optimal snit-GFP/YFP nanobodies for this task. Additionally, a YFP (or GFP) tag may not always be suitable, for example in Sf-9 Easy Titer cells, which express GFP upon infection with baculovirus (16). Furthermore, a second, orthologous tag that enables simultaneous monitoring of multiple protein constructs or sequential affinity purifications would be very beneficial for protein purification.

We sought to determine whether mCherry, another widely used fluorescent protein with fluorescence excitation and emission spectra that are distinct from that of YFP (17) could be used to generate an affinity resin suitable for large-scale recombinant protein production. Since nanobodies that recognize mCherry have also been described (18), we evaluated three candidate anti-mCherry nanobodies alongside three anti-YFP/GFP nanobodies for specific criteria to determine whether they make a reliable and scalable recombinant protein affinity support, and which nanobodies performed the best at this task. We evaluated yield and purity of nanobodies expressed in *E. coli*, as well as the specificity, binding capacity, and re-usability of supports generated using these nanobodies. We used X-ray crystallography to define the molecular basis of GFP/mCherry binding that underlies their utility as an affinity matrix. Finally, we demonstrated that the mCherry-Tag system, comprised of anti-mCherry nanobody-sepharose and N-terminal fusion of mCherry to a protein of interest can yield milligram-scale quantities of recombinant Topoisomerase 2.

## Results

### Generation of anti YFP/mCherry affinity matrices

We sought to develop a robust affinity resin with binding characteristics comparable to the YFP-tag (9) and would not cross-react with the YFP tag. Therefore, we examined the utility of an affinity resin that binds to a red fluorescent protein. mCherry (19) is derived from the dsRed protein of *Discosoma sp*. and shares low sequence identity (<30%) to GFP from *Aequorea victoria* and its commonly used variants, which includes YFP. A report from the Rout group (18) described a panel of camelid atypical immunoglobulins, commonly referred to as nanobodies that recognize mCherry or GFP. We selected from these three llama mCherry (LaM) and two GFP/YFP (LaG) nanobodies with binding affinities (Supplementary Table 1) comparable to that of the GFP-enhancer nanobody used with the YFP-tag affinity support (9,12) and evaluated their use as a recombinant protein purification platform.

To determine the suitability of each nanobody for large-scale production of an affinity matrix, we first evaluated the purity and yield of each nanobody expressed in *E. coli* with an N-terminal His_6_–tag. After Ni-NTA purification, we analyzed the crude nanobody protein using size-exclusion chromatography (Figure 1). Each of the nanobodies eluted as a monodisperse peak that was separate from protein aggregates and contaminants. Most of the nanobodies eluted from the column near the volume expected for the monomeric protein of 82 mL, corresponding to an apparent mass of 15 kDa for the His_6_-tagged protein. Interestingly two of the nanobodies (GFP-enhancer and LaM2) eluted about 10 mL later than expected. This could be due to weak interactions with the sepharose matrix rather than a smaller size, since these nanobodies elute even later than Ubiquitin, a much smaller protein (8.5 kDa).

**Fig. 1.**
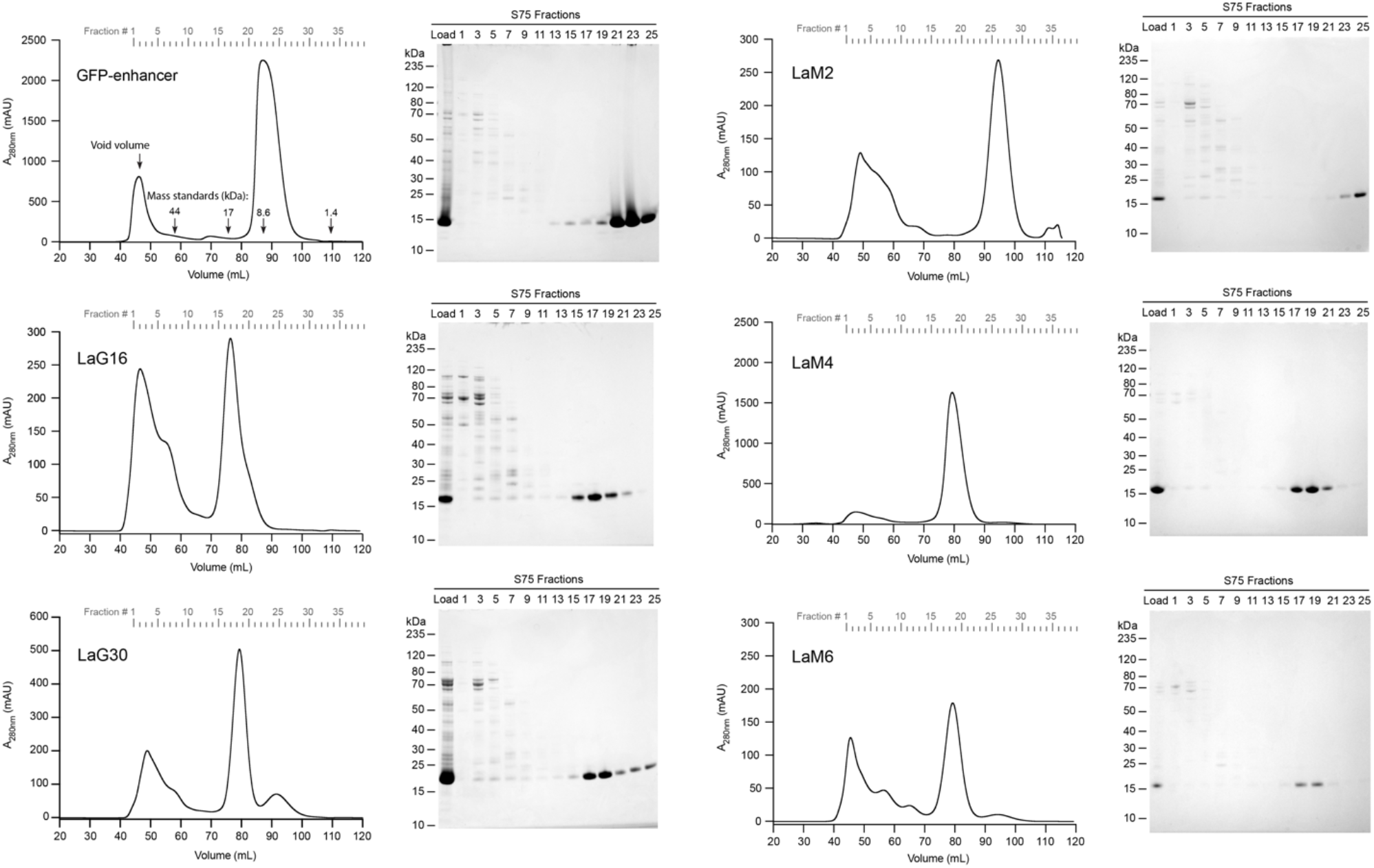
Purification of nanobodies. Representative FPLC chromatograms and SDS-PAGE gels of the indicated anti-GFP or anti-mCherry nanobodies purified using the size exclusion S75 column. Samples of column load and the indicated elution fractions that span the size range from aggregates to monomeric nanobody peaks in each chromatogram are shown on the corresponding Coomassie blue stained SDS-PAGE gel.

From the FPLC chromatograms, expression yield of each nanobody in *E. coli* can be estimated from the area of the A_280_ absorbance peak in the chromatogram. Between the three anti-GFP nanobodies that we studied (Figure 1A, 1B & 1C), the GFP-enhancer nanobody yielded much more protein than LaG16 and LaG30 and had the lowest amount of aggregated protein, which elutes near the column void volume. Among the three anti-mCherry nanobodies, LaM4 had the largest monomer peak area and therefore protein yield, with a lower proportion of contaminants and protein aggregates than LaM2 or LaM6 (Figure 1D, 1E & 1F). Fractions containing pure nanobody were pooled and quantified (Supplementary Table 2). Yields of about 30-60 mg purified nanobody per liter of culture were obtained for LaM4 and GFP-enhancer, which is 3 to 10-fold higher than the others, indicating that nanobody yield can be quite variable and is an important parameter to evaluate.

We examined whether the differences in yield also correlated with protein stability as measured using a protein thermal shift assay (Supplementary Table 3 and Supplementary Figure 1). In this assay nanobodies are incubated with Sypro Orange dye that fluoresces when it binds to hydrophobic residues that are exposed upon heat-induced denaturation, with the midpoint of the fluorescence change defining the melting temperature (T_m_) of the protein. LaM2 had the highest T_m_, despite the significantly lower recombinant protein yield compared to LaM4 or GFP-E nanobodies. The other three low-expressing nanobodies (LaG16, LaG30, and LaM6) had lower T_m_ values suggesting that although a high T_m_ is not necessarily an indicator of a high yield, a low T_m_ may indicate that a nanobody is less likely to express at a high level.

### Molecular basis of nanobody:FP interactions

To determine the molecular architecture of the nanobody:FP interactions, we purified 1:1 complexes with their cognate FP and performed crystallization trials. We obtained diffraction-quality crystals for LaG16, LaG30, LaM2, LaM4, and LaM6 nanobody-FP complexes and solved crystal structures for each at resolutions ranging from 1.15Å to 2.37Å (Figure 2, Supplementary Table 4). The structure of the GFP-enhancer nanobody bound to GFP has previously been described (12). The nanobodies consist of the canonical Ig fold that contains antiparallel beta sheets with a few short alpha helices at each end. eGFP and mCherry both have the expected beta barrel shape. Each nanobody bound its cognate FP through an interface that buried between 520 and 730Å^2^, which is an interface size typical of nanobodies with nanomolar affinity for their antigen (20). Among the anti-GFP nanobodies, LaG16 and LaG30 nanobodies adopted the same binding positions with eGFP (RMSD 0.35Å), which is not unexpected since they share 90% sequence identity. The difference between the binding positions on the anti-mCherry nanobodies is more distinct. LaM4 sits directly above one end of the β-barrel. LaM2 and LaM6 bind the β-barrel surface near one edge of mCherry and share a binding epitope, but they bind through distinct molecular interactions. We examined the locations of lysine ε-amino groups (depicted as spheres in figure 2) to evaluate whether immobilization of the nanobody using chemical crosslinking to NHS-sepharose could interfere with the nanobodies high affinity FP binding-site. We did not observe any lysines within hydrogen-bonding range of the FP binding epitope, indicating that covalent coupling to NHS-activated sepharose to generate an affinity matrix is unlikely to alter the FP binding epitope.

**Fig. 2.**
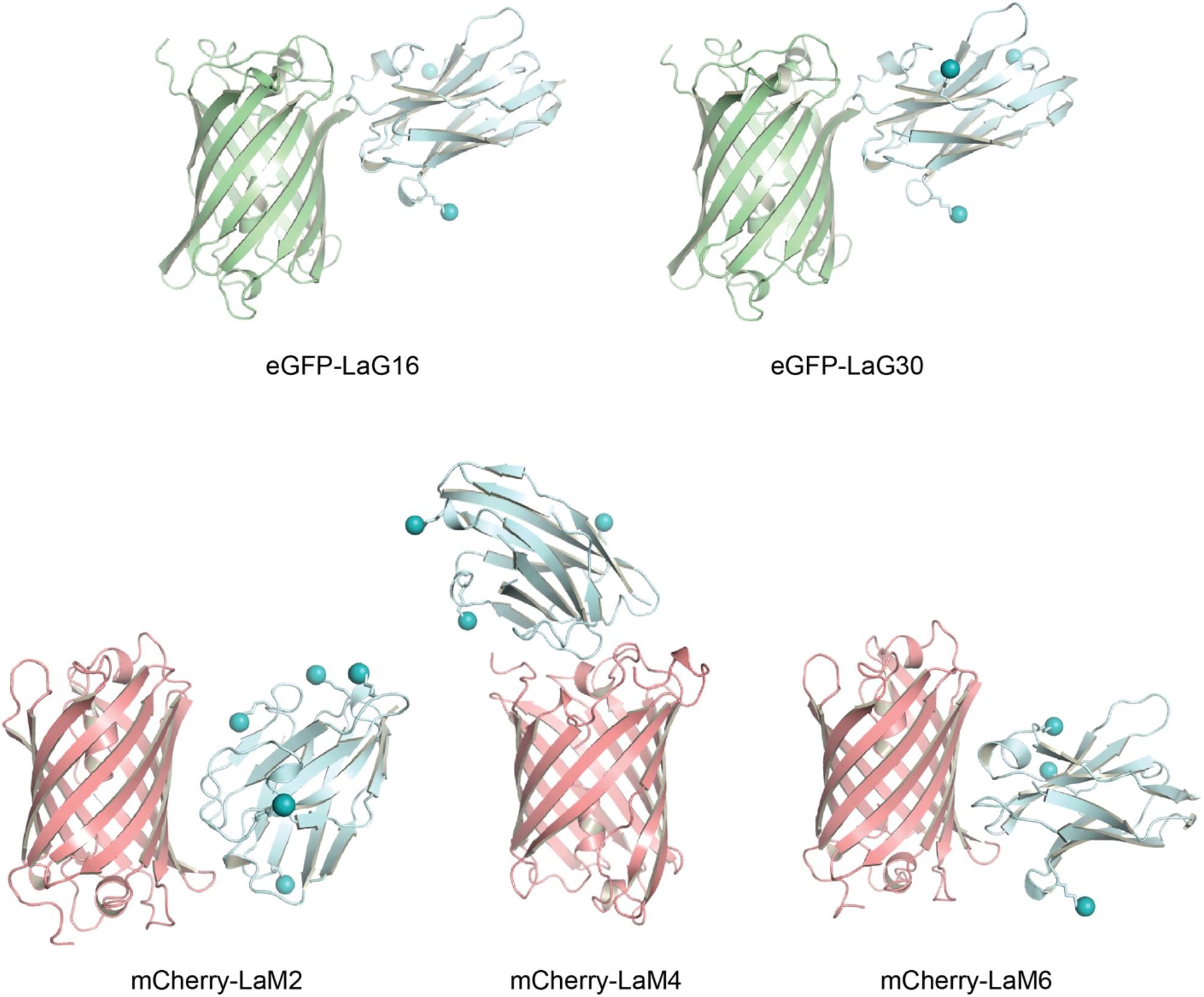
Structures of Nanobody:FP complexes. Cartoon depiction of X-ray crystal structures of the indicated nanobodies bound to their cognate FP. Spheres indicate the location of ε-amino group of lysine residues.

### Selectivity and stability of the nanobody affinity matrices

We coupled an equal quantity of each nanobody to NHS-activated sepharose to generate an affinity support and evaluated binding capacity and specificity of these resins. We challenged each resin with a solution containing both enhanced GFP (eGFP) and the monomeric red fluorescent protein mCherry in equal quantities sufficient to saturate the resin (Figure 3A). After a one-hour incubation we could visually observe binding of the correct fluorescent protein as a change in resin color. Each of the anti-GFP and anti-mCherry nanobody resins bound their cognate FP and became brightly colored, leaving the supernatant colored predominantly by the unbound FP. An SDS-PAGE analysis of protein eluted from the nanobody resin showed that each nanobody bound similar amounts of its cognate FP, but with LaG30 and LaM2 slightly under-performing their competitors. They are also specific for their target FP as no eGFP was bound by LaM2, LaM4, or LaM6, while no mCherry bound to GFP-enhancer, LaG16, or LaG30. We note that a portion of the purified untagged mCherry protein used in our experiments retains some structure in an SDS-PAGE gel, resulting in an anomalously migrating band. This band is likely degraded mCherry and is only present in our mCherry protein sample (Supplementary Figure 2).

**Fig. 3.**
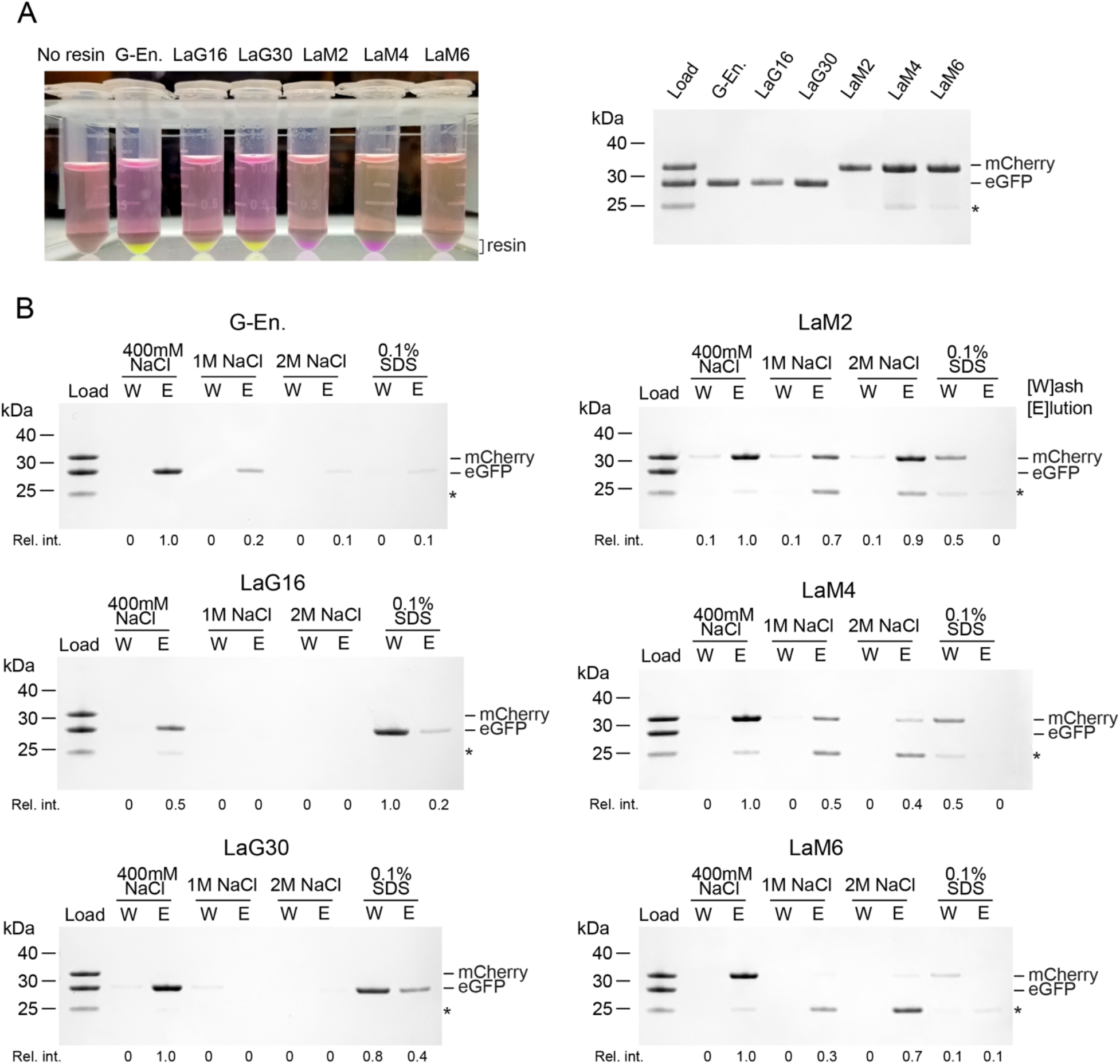
Binding tests and buffer compatibility of nanobody-sepharose resins. (A) The indicated nanobody-sepharose binding reactions (left) with resin bound to saturating amounts of the target FP visible at the bottom of the tube. G-En is GFP-enhancer nanobody. (right) SDS-PAGE of bound protein shows that binding can be monitored visually, and no off-target binding is observed. * Indicates band corresponding to degraded mCherry. (B) Indicated nanobody-sepharose resins were incubated with saturating amounts of eGFP and mCherry, then washed 3 times with the indicated buffers. The last wash is shown (W) and protein that remained on the resin and was eluted from the column with 0.1M glycine pH 2 is shown (E) analyzed by Coomassie blue-stained SDS-PAGE. Relative intensity of bound FP is indicated below each lane.

We evaluated the ability of the nanobody resins to retain eGFP or mCherry under more stringent wash conditions such as high salt or the detergent SDS (Figure 3B). After three washes of 10x the resin volume, the GFP-enhancer and LaM4 nanobodies retained 20–50% of the bound FP at 1M NaCl and less so at 2M NaCl, suggesting the maximum NaCl should be kept below 1M NaCl to avoid significant protein loss. LaG16 and LaG30 did not retain FP in 1M NaCl, suggesting these nanobodies are highly sensitive to salt. LaM2 retained mCherry even at 2M salt, however, we note that the presence of some mCherry in each of the washes suggests that even though the LaM2-mCherry interaction is quite stable to salt, mCherry will slowly dissociate during washing. We also examined the ability of the nanobody resins to retain FP after washing with 0.1% SDS. Most, but not all proteins will denature under this concentration of SDS, making such conditions useful when the aim is to avoid unwanted enzyme activities such as removal of post-translational modifications during purification. Although most of the nanobody-FP interactions were disrupted by 0.1% SDS, LaG30 retained some eGFP suggesting that it is the most stable to SDS amongst our panel. LaG16 and GFP-enhancer retained only trace amounts and the mCherry nanobodies retained little to no FP, which indicates they perform poorly in the presence of the denaturant SDS.

### Performance of nanobody affinity matrices

We used lysates from *E. coli* or HEK293F cells to challenge the nanobody resins with a complex protein mixture (cell lysate) containing eGFP/YFP or mCherry. First, we performed binding tests using *E. coli* and HEK293F cell lysates that contained fluorescent proteins to evaluate each resin (Figure 4). Equal amounts of each resin were incubated with each of the cell lysates: *E. coli* expressing mCherry, *E. coli* expressing eGFP, HEK293F expressing mCherry-TOP2α, and HEK293F expressing YFP-TOP2α. The final eluates were run on an SDS-PAGE gel and stained with Coomassie blue to determine the protein composition present in each eluate. All six resins exhibited good selectivity against their non-target fluorescent proteins while enriching their target fluorescent proteins. Importantly, in the absence of cognate FP, the eluate is devoid of detectable proteins, indicating that affinity resin generated from FPLC-purified nanobodies exhibits very low off-target binding. This observation held true for not only proteins expressed in *E. coli* cells (Figure 4A) but also HEK293F cells (Figure 4B). The amount of fluorescent protein isolated by each resin, shown by the intensity of the eluate bands, varies somewhat amongst the nanobodies. While all the resins were significantly enriched with their target FP amongst the GFP/YFP nanobodies, the GFP-enhancer resin bound slightly more FP from both lysates. LaM4 resin bound more mCherry in *E coli* lysate, while LaM2 appeared to bind more mCherry-TOP2α from HEK293F lysate. This suggests that while all the tested nanobodies can perform as affinity matrices, there is a difference in binding capacity between these nanobody resins, with LaG16, LaG30, and LaM6 slightly underperforming the others.

**Fig. 4.**
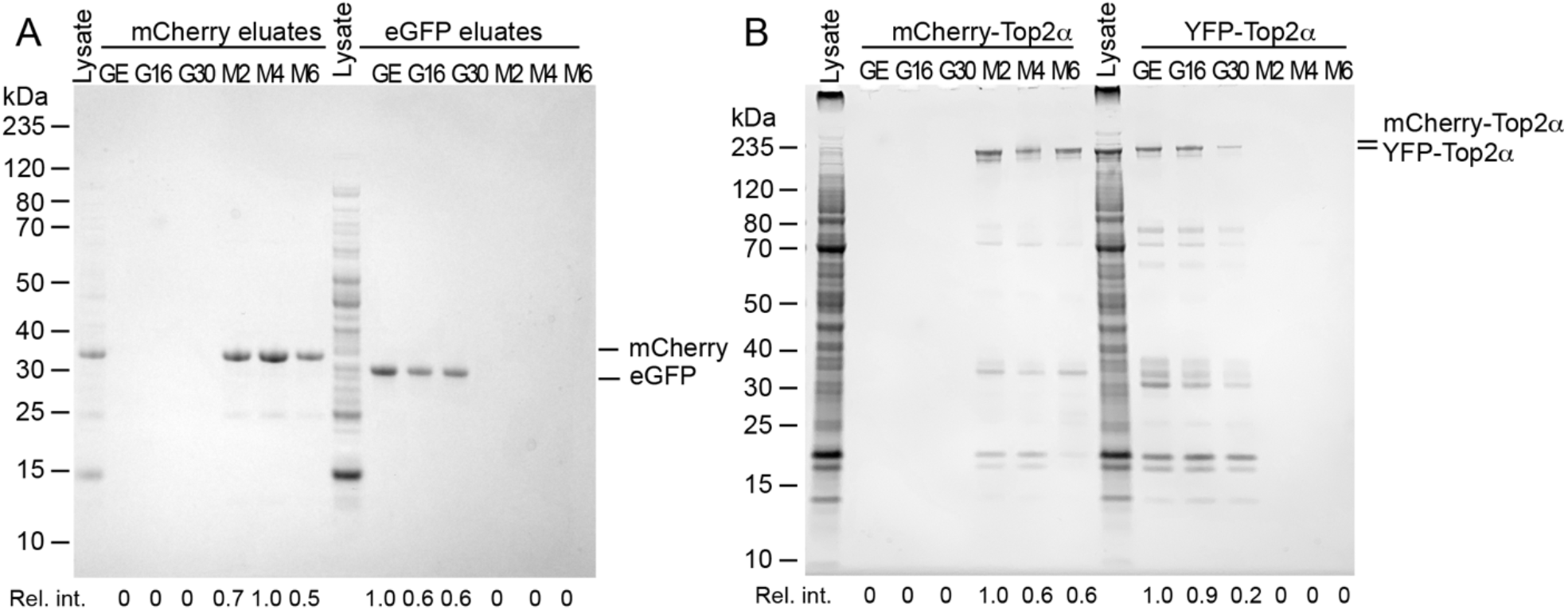
Evaluation of nanobody efficiency and specificity using cell lysates. All six nanobody-sepharose resins including GFP-enhancer (GE), LaG16 (G16), LaG30 (G30), LaM2 (M2), LaM4 (M4), and LaM6 (M6) were incubated in the same amount of cell lysate, then bound protein was eluted with 0.1 M glycine pH 2. (A) *E. coli* cell lysates that express mCherry or eGFP were incubated with the indicated nanobody-sepharose resins. Coomassie blue stained SDS-PAGE of eluates is shown. (B) Lysates from HEK293F cells transfected with mCherry-TOP2α or YFP-TOP2α were incubated with the indicated nanobody-sepharose resins. Coomassie blue stained SDS-PAGE of eluates is shown. Relative intensity of bound FP is indicated below each lane.

### Stability and Reusability of nanobody-sepharose affinity matrices

Given the cost and effort associated with preparing a nanobody-sepharose affinity resin for recombinant protein production, it is very important that such a resin be re-useable. Since bound FP cannot be removed easily, harsh removal conditions such as 0.1M glycine pH 2 are required to disrupt the nanobody–FP interaction and regenerate the FP-binding sites. Since low pH could potentially denature the nanobodies as well, we performed 20 cycles of binding saturating quantities of their cognate FP to each nanobody-sepharose resin, followed by washing and elution with pH 2 glycine buffer to evaluate how the nanobody resins would retain their binding capacities after many rounds of recombinant protein purification. We compared the resins in the first binding cycle (Figure 3A) vs the 20^th^ binding cycle (Figure 5B) and observed both by the color of bound FP and SDS-PAGE analysis of the eluate that there was no detectible decrease in resin capacity for GFP-enhancer, LaM2, and LaM4, suggesting that their useable lifetime will extend well beyond the 20 rounds assayed. LaG16, LaG30, and LaM6 again displayed reduced capacity suggesting they are not stable to repeated cycles of binding and regeneration. To determine the stability of resins at pH2 specifically, we evaluated the stability of each nanobody sepharose after 16 hours incubation at room temperature in glycine pH 2 (Figure 5C). A comparison to new resin (Figure 3A) shows that GFP-enhancer, LaM2, and LaM4 nanobody resins have retained their binding capacity, as demonstrated by SDS-PAGE analysis of eluates. LaG16, LaG30, and LaM6 have reduced binding capacity suggesting the pH 2 regeneration step is causing the nanobody to denature, resulting in reduced resin binding capacity. We note that these same 3 nanobodies had the lowest melting temperature (Supplementary Figure 1), suggesting that a low T_m_ is an indicator of sensitivity to low pH regeneration. A higher T_m_ indicates greater protein stability and a lower free energy of protein folding. Because harsh conditions like that of pH 2 destabilize protein structure, the folded state is more likely to persist for proteins with a higher T_m_.

**Fig. 5.**
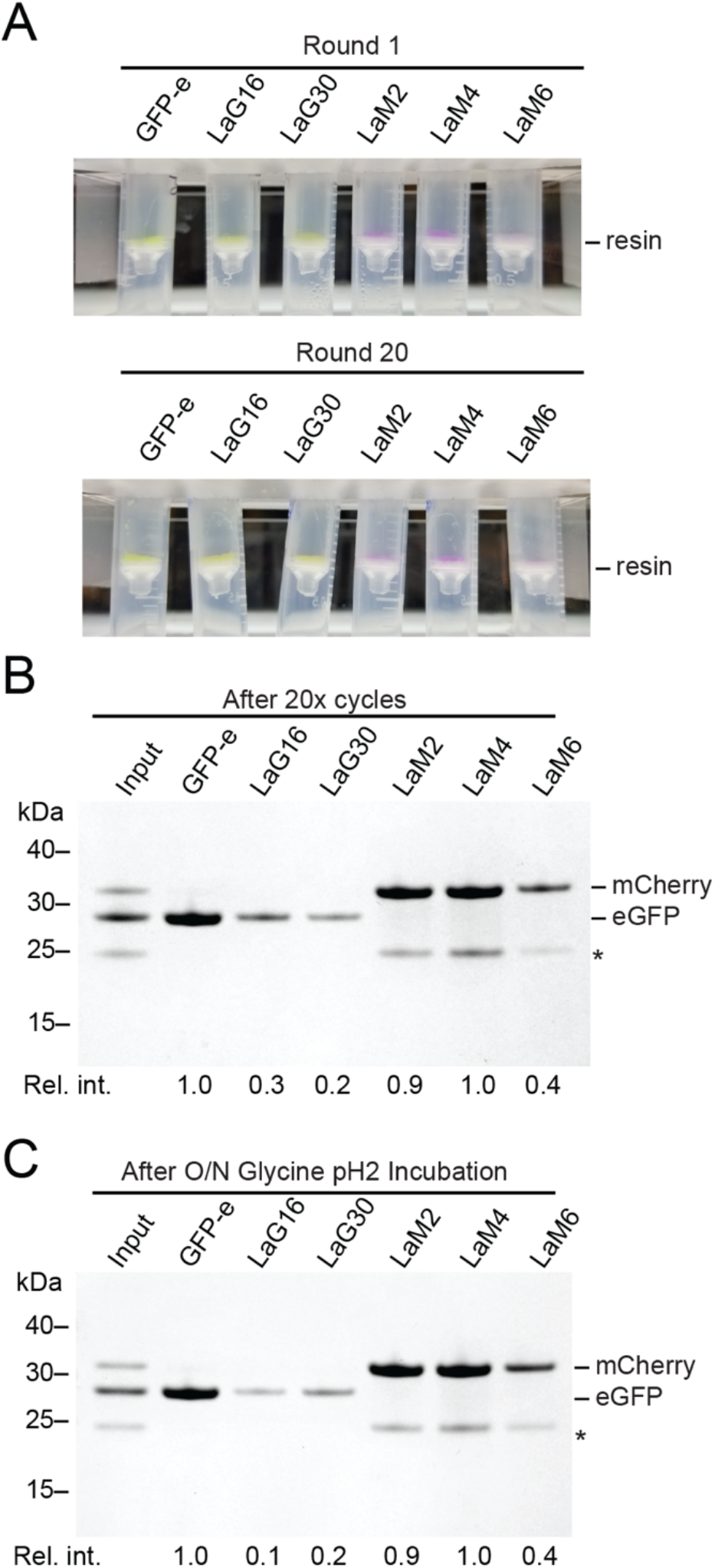
Stability of nanobody sepharose resins to regeneration at low pH. (A) Nanobody-sepharose resins after binding to saturating amounts of FP in the initial round, or after 20 cycles of binding and regeneration. Binding of FP to resin can be observed visually on the spin filter where the resin accumulates. GFP-e is GFP-enhancer nanobody. (B) SDS-PAGE of protein bound to resin in panel A after 20 cycles of binding and regeneration. This is comparable to the first cycle of binding as shown in figure 3A. * Indicates band corresponding to degraded mCherry. (C) SDS-PAGE analysis of protein bound to nanobody-sepharose resins that had been pre-incubated in glycine pH 2 for overnight. Relative intensity of bound FP is indicated below each lane.

### Purification of recombinant TOP2α using LaM4 affinity resin and an mCherry-tag

Our goal is to establish an anti-mCherry affinity purification system that is suitable for large-scale protein purification of mCherry-tagged proteins with comparable efficiency to the GFP/YFP-tag system. Our overall expression strategy is outlined in Figure 6. First, a plasmid DNA expression construct is generated which encodes the protein of interest fused to a fluorescent protein tag with an intervening protease cleavage site. These plasmids are transiently transfected into a suspension culture of HEK293F cells using polyethylenimine (PEI). Protein expression occurs over the course of 48-72 hours, during which the level of protein expression in transfected cells can be monitored via the fluorescent signal from mCherry. Recombinant mCherry-tagged protein is isolated from cell lysate by binding to LaM4-sepharose, then eluted by protease cleavage for further purification and polishing by FPLC. The resin can be regenerated then reused by washing with Glycine pH 2.

**Fig. 6.**
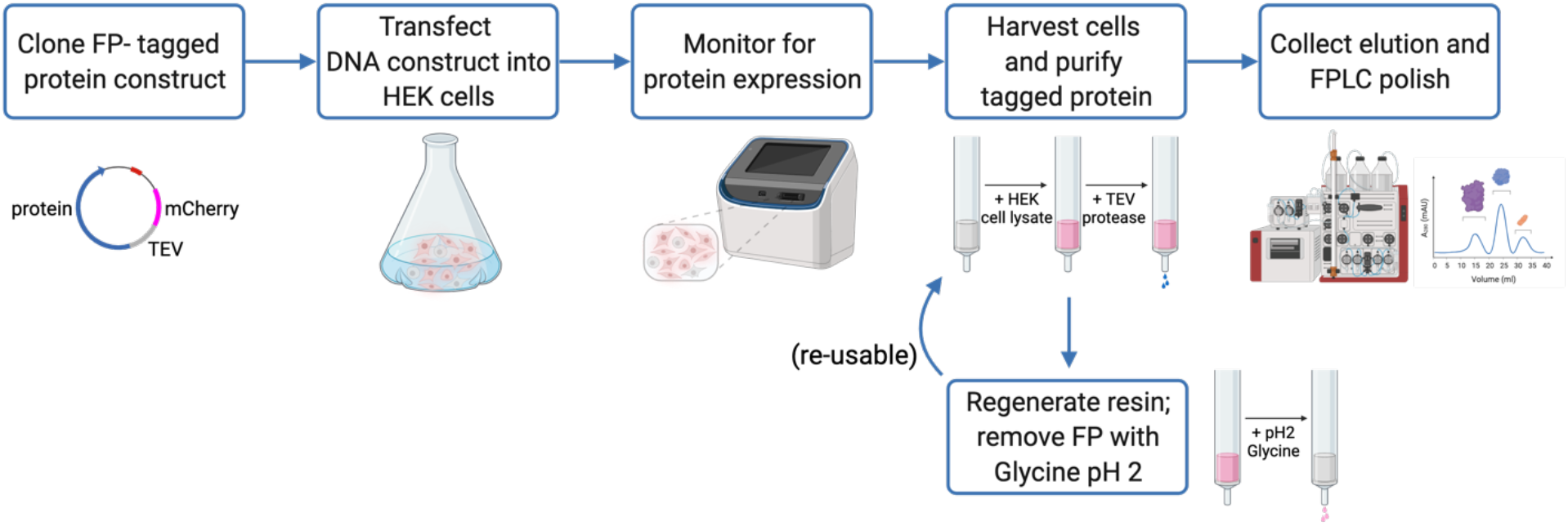
Flow diagram of nanobody-based affinity purification system. Plasmid DNA encoding the protein of interest fused to a fluorescent protein tag with an intervening protease cleavage site is transfected into a culture of HEK293 cells in suspension. The recombinant protein fluorescent signal can be monitored during the expression time course. Cells are grown and monitored for fluorescent protein expression signal over the course of <u>48-</u>72 hours before lysis. Lysate containing FP-tagged protein is applied over a nanobody resin, and the protein of interest is eluted by protease cleavage for further FPLC polish. FP-bound resin can be regenerated by washing with Glycine pH 2 solution and reused for future protein purification.

We proceeded to challenge LaM4 nanobody resin by testing its efficacy in a large-scale protein purification. Type 2 Topoisomerases are important biological enzymes, and assays that use recombinant human TOP2 are important for evaluating its enzymatic activity as well as the effects of anti-cancer chemotherapeutic drugs. The YFP-tag system enables high-yield (10-25 mg per liter of culture) (9) expression and purification of full-length recombinant TOP2α from suspension-cultured HEK293F cells. To evaluate the performance of the mCherry-tag system, we generated an expression vector that encodes mCherry-TOP2α and followed the same transfection and purification protocol as was used for YFP-TOP2α. The GFP-enhancer resin was substituted with LaM4 resin for protein purification. The dark magenta color observed in the cell pellet (Figure 7A) was indicative of robust mCherry-TOP2α expression (Figure 7B), and mCherry-TOP2α binding to the LaM4 resin could be observed as a change in resin color (Figure 7C). TEV protease cleavage releases untagged TOP2α with an N-terminus that corresponds to the native protein (Figure 7D), and it could be purified to homogeneity by sequential ion exchange (Figure 7E) and size-exclusion chromatography (Figure 7F) to yield protein in comparable purity to that obtained using the YFP-tag system. A 0.5L culture yielded 2.6 mg of purified TOP2α, for a yield of 5.2 mg/L which is comparable, although slightly lower than the 10 mg/L yield reported for the YFP-tag (9).

**Fig. 7.**
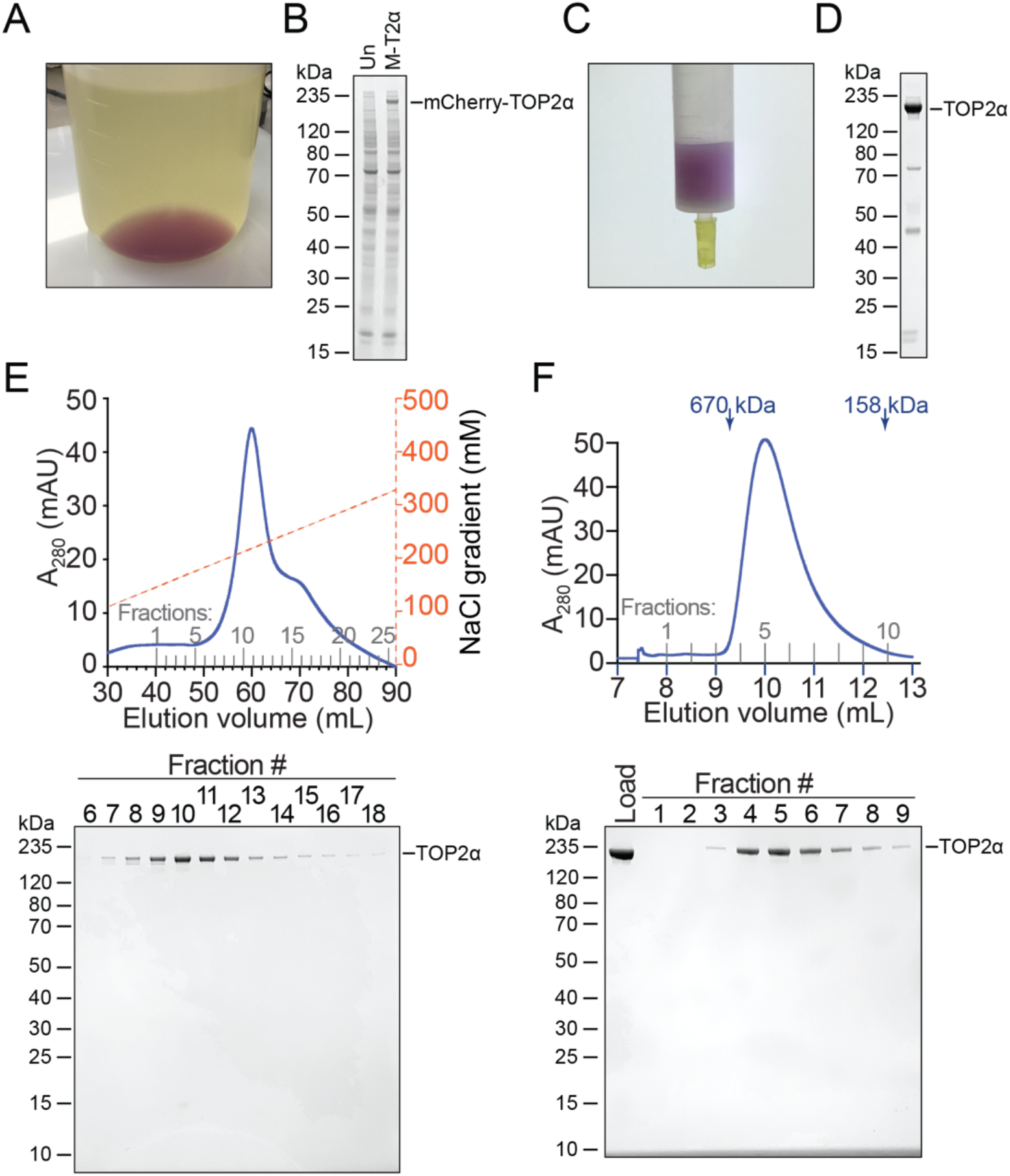
Purification of Topoisomerase 2α using LaM4-sepharose and an mCherry tag system. (A) HEK293F cells were transfected with plasmid encoding mCherry-TOP2α and pelleted by centrifugation after 3 days. A dark purple color indicates robust expression of the mCherry-tagged protein. (B) Coomassie blue-stained SDS-PAGE of untransfected HEK293F cells (Un) or HEK293F cells transfected with mCherry-TOP2α (M-T2α) showing robust expression of mCherry-TOP2α visible as a band in total cellular lysate. (C) Binding of mCherry-TOP2α to LaM4-sepharose can be monitored visually. (D) SDS-PAGE of TEV-eluted TOP2α. (E) FPLC cation exchange purification of TOP2α and SDS-PAGE of the indicated fractions. (F) FPLC size exclusion chromatography purification of TOP2α and SDS-PAGE of the indicated fractions.

## Discussion

Here we evaluate a panel of nanobodies that bind YFP/GFP or mCherry for their use as an affinity matrix for recombinant protein production. All 6 nanobodies, when coupled to NHS-activated sepharose functioned as efficient affinity matrices that strongly bound their target FP and not to non-target FP. This high degree of specificity makes dual-tagging systems possible, where sequential purification steps can be used to purify co-expressed proteins. It is important to evaluate nanobodies using multiple parameters when identifying a suitable candidate to use for an affinity matrix. Amongst the nanobodies assayed here, the GFP-Enhancer and LaM4 nanobodies produced affinity resins with the highest capacity, albeit only slightly higher than the other nanobodies. Notably, GFP-Enhancer and LaM4 nanobodies have a significant advantage of 5 to 10–fold higher expression yield from the *E. coli* culture that is used to generate the nanobodies and are therefore the optimal choice for preparing large batches of affinity matrix. LaM2 is stable to high salt (up to 2M NaCl) but also slowly releases mCherry during washing which will ultimately reduce the yield. LaG16, LaG30, and LaM6 are sensitive to salt concentrations, yet this could be considered an advantage when it is desirable to elute the FP with high salt rather than with a site-specific protease. Reusability of affinity matrix is important when used on a large scale for recombinant protein production, and we have found that the protein melting temperature is an important predictor of stability to low pH used for resin regeneration. In our hands we have not been able to identify a suitable alternative resin wash condition, as other mild denaturants such as 0.1% SDS or high NaCl fail to remove all the FP (Figure 3B). Therefore, a nanobody sepharose resin must be stable to washing at low pH if it is to be reusable. A recent report described the use of a tethered GFP-enhancer–LaG16 nanobody pair as an affinity matrix with improved affinity for GFP (21). However, our finding that LaG16 is not stable to multiple cycles of regeneration and re-use suggests that such a resin would have a limited useful lifetime. Another recent report identified a nanobody called Nb2 that bound to the same epitope as LaG16 and was more thermostable (22), suggesting a GFP enhancer-Nb2 nanobody tandem fusion could generate an affinity matrix that could survive cycles of regeneration and reuse.

All the nanobodies assayed exhibited a high specificity for their antigen and we did not observe any off-target binding, yet there are clear differences in their performance characteristics. Considering all the parameters on which the affinity matrices were evaluated, we have selected GFP-Enhancer- and LaM4-sepharose as the optimal pair for recombinant protein production. We now routinely use GFP or YFP affinity tags paired with GFP-enhancer nanobody sepharose, and mCherry tags paired with the LaM4 nanobody sepharose for recombinant protein production. For both YFP and GFP, the nanobody binding epitope residues are the same, and therefore the GFP-enhancer affinity matrix is equally suitable for purifying GFP or YFP-tagged proteins. The nanobody epitopes contain neither the N- or C-terminal residues of the FPs, which indicates that FPs should perform equally well as either an N-terminal or C-terminal affinity tag. These tags confer specific advantages when expressing recombinant protein in HEK293F cells. Overexpression of FP-tagged proteins produces a very high fluorescence signal that can be conveniently monitored in real time. Regardless of the type of media the fluorescence signals from cellular expression are vastly greater than the background media fluorescence. In addition, we can also visualize quantity of protein binding to their cognate affinity matrix. These features allow usto troubleshoot and optimize transfection conditions, growth time, buffer conditions, and binding to resin. In contrast, non-fluorescent tags require time-consuming procedures such as western blotting or SDS-PAGE to accomplish the same task. Thus, we present the GFP/YFP and mCherry tags as two cost-effective and efficient affinity purification systems that overcome the challenges associated with recombinant protein expression in cultured mammalian cells. There is a vast array of expression vectors that can be used to express proteins tagged with a FP, and expression of FP-tagged proteins in mammalian cells is a widely used technique for microscopy. Plasmid repositories have plasmids already available that express many proteins as FP-fusions for use in live-cell imaging or microscopy experiments. There is a diverse array of resources available for generating FP-tagged expression constructs, which lowers the barrier to initiating an FP-based affinity purification.

We have described X-ray crystal structures of the nanobody-FP complexes that form on the affinity resins used in this work. NHS-activated sepharose is commonly used to generate protein-linked sepharose due to its cost, chemical compatibility, and reactivity with the primary amines of lysine residues or the N-terminus. Since NHS esters acylate the lysine side-chains of a protein, it is important that lysines are not critical components of the binding interface. Our structures reveal there are no nanobody lysine residues at the FP-binding site. Chemically linking the nanobody to NHS-activated sepharose should not alter the FP binding site, and indeed all the nanobody sepharose resins bound their target FP (Figure 3A). Furthermore, our structures provide a means by which to predict whether one of the many FP variants (17) will bind to a given nanobody prior to experimental testing. Finally, these structures add to the repertoire of nanobody structures, which will aid in the future development of synthetic nanobodies (23), which can produce high-affinity binding molecules without the use of animals.

### Experimental Procedures

#### Nanobody expression in E. coli

Synthetic DNA constructs encoding LaM2, LaM4, LaM6, LaG16, and LaG30 nanobodies(18) optimized for expression in *E*. coli (Genscript) were cloned into the NdeI and XhoI sites of pET-15d (Novagen). Plasmid encoding GFP-enhancer nanobody has been previously described (9). Each plasmid was transformed into Rosetta2 cells (EMD) and plated on LB-agar media with 100 μg mL^-1^ ampicillin and 34 μg mL^-1^ chloramphenicol. 10 mL cultures were inoculated and grown overnight at 37°C in the same antibiotics. For protein expression, cultures were prepared by transferring the starting cultures to 2L bottles of Terrific Broth media with 250 μL of Antifoam204 (Sigma), then incubated at 37°C in a Lex-48 Bioreactor (Ephiphyte3) until cultures reached an OD600 value near 2. To each 1 L culture, 50 uL of 1 M IPTG was added and the cells were further incubated at 16°C overnight. *E. coli* cultures were spun down in a Lynx 4000 centrifuge (Thermo Fisher) at 6,000 x g for 20 minutes, and cell pellets were transferred to 50 mL Falcon tubes and stored at -80°C until ready for protein extraction.

#### Purification of Recombinant Nanobodies

Cell pellets frozen at -80°C were thawed in a water bath and resuspended in lysis buffer (20 mM Tris pH 7.5, 300 mM NaCl, 10 mM imidazole, and 0.5 mM TCEP), then transferred to a chilled metal beaker. 10 mg of lysozyme (Goldbio) and 50 μL of saturated PMSF (Goldbio) in ethanol were added, and the mixture was incubated on ice for 30 minutes with occasional mixing. Cell lysate was sonicated at 80% power in three 30 second intervals, with 1-minute cooling in between to reduce the viscosity. Clarified lysate was collected after centrifugation at 25,000 g for 30 minutes and passed over a Ni-NTA resin bed that was equilibrated in lysis buffer. After washing six times with running buffer to remove remaining cell debris or unbound proteins, nanobodies were eluted using running buffer supplemented with 250 mM imidazole. Elution fractions were analyzed by SDS-PAGE, and fractions containing the nanobody protein were pooled together and precipitated with 2 volumes of 4 M ammonium sulfate followed by centrifugation at 25,000 x g for 30 minutes at 4 °C and kept at -80°C until further purification using FPLC.

Frozen protein pellets were thawed and re-dissolved by addition of a minimal amount of Milli-Q water, then centrifuged at 4°C, 20,000 x g for 10 minutes to eliminate any insoluble particulates. Clarified solution was injected into an ÄKTA go FPLC system (Cytiva) and purified on a HiLoad 16/600 Superdex 75 pg column (Cytiva) using 1X PBS buffer at a flow rate of 1 mL min^-1^. The column was calibrated using Gel Filtration Standards (Bio-Rad) and recombinant ubiquitin protein (24) run in the same PBS buffer. Elution for each batch of nanobody was collected in fractions of 2 mL volume, tested on a Coomassie blue stained SDS-PAGE. Fractions containing nanobodies were pooled together and concentrated using 10K cut off centrifugal filters (Millipore).

#### Preparation of nanobody sepharose

Nanobody sepharose was prepared as described previously (9). Briefly, each purified nanobody at 10 mg mL^-1^ concentration was coupled to NHS-activated Sepharose 4 fast flow (Cytiva) at a ratio of 2:1 nanobody solution to resin slurry. This mixture of resin and nanobody was incubated on a nutator overnight at 4°C. Crosslinking efficiency was verified by the absence of protein in the supernatant when measured using a Bradford assay (Biorad). Resin was then quenched with 100 mM Tris pH 8.0, 5 mM BME at room temperature for 1 hour. Quenched resin was washed with two wash buffers (buffer 1: 100 mM Tris pH 9, 500 mM NaCl; buffer 2: 100 mM NaOAc pH4.5, 500 mM NaCl) alternating for six cycles total. Nanobody resins were equilibrated in storage buffer (50 mM Tris pH 8.0, 400 mM NaCl, 0.1% NP-40) and stored at 4°C until ready for use.

#### Characterizations of nanobody sepharose

Prepared nanobody resins were tested with lysates of *E*. coli that were expressing fluorescent proteins (eGFP or mCherry) or lysates of HEK293F cell lysate that expressed either YFP-Top2α (9) or RFP-Top2α. For each lysate, a mixture including 425 μL storage buffer (50 mM Tris pH 8.0, 400 mM NaCl, 0.1% NP-40), 25 μL cell lysate, and 50 μL resin slurry was prepared for each resin, followed by a 30-minute incubation on a nutator at 4°C. The mixtures were then transferred into disposable spin columns and centrifuged at 1000 g for 10 seconds. The flow-through was discarded and resins were washed with 400 μL of storage buffer three times. Fluorescent proteins were eluted from the resins using 2 volumes of 25 μL 0.1 M Glycine pH 2. All six resins were tested against lysates that expressed eGFP or mCherry individually. Elution from each nanobody resin was analyzed by SDS-PAGE stained with Coomassie blue to compare their binding ability and selectivity.

To evaluate the binding capacity of nanobody-sepharose resin, 25 μL of resin was incubated with 15 μM eGFP and 15 μM mCherry in 1 mL of storage buffer. Slurries were set on a nutator at 4°C for 2 hours, and again transferred into new disposable spin columns to centrifuge at 1000 g for 10 seconds, washed three times with 400 μL storage buffer (with additional NaCl or SDS where indicated), then protein was eluted with three volumes of 25 μL 0.1 M Glycine pH 2.

Equal volumes of elution from each resin were tested side by side on a Coomassie blue stained SDS-PAGE gel to evaluate the amount of FP bound by each resin. Gels were photographed and band intensities were quantified using FIJI (ImageJ; NIH) with background intensity measured using an identical sized box immediately above or below each band subtracted. The intensity of the anomalously migrating mCherry band marked as a * in each gel was included in the calculation for the amount of mCherry present. The lane with the highest amount of FP was defined as an intensity of 1, and weaker numbers as a fraction thereof.

#### Crystallization of purified nanobody-FP

FPLC-purified nanobodies were concentrated to 17 mg/mL using a spin concentrator (Millipore) and incubated in a 1:1 molar ratio with their respective fluorescent protein binding partner for 30 minutes at 4°C. Crystal of nanobody-fluorescent protein complexes were grown using the sitting-drop vapor diffusion method by mixing 200nL of protein complex mixture and 200nL of precipitant. The reservoir contained 25 µL of each crystallization solution. Crystals of GFP-LaG16 complex were obtained in 100mM BICINE pH 9.0 with 2% (w/v) 1,4-Dioxane and 10% (w/v) PEG20000. For the GFP-LaG30 complex, crystals were obtained in 800mM monopotassium phosphate, 800mM monosodium phosphate, and 100mM HEPES pH 7.7. These crystals were transferred into cryoprotectant solution containing 100mM Tris pH 9.0, 10% (w/v) PEG6000, and 25% (w/v) glycerol for GFP-LaG16 and crystallization condition supplemented with 25% (w/v) glycerol for GFP-LaG30. mCherry-LaM2 crystals were obtained in 2.5M malonate pH 7.0 and transferred into 3M malonate pH 7.0 with 15% (w/v) glycerol for cryoprotection. Crystals of mCherry-LaM4 complex were obtained in 100mM MMT (DL-Malic acid, MES, and Tris in a 1:2:2 molar ratio) pH 4.0 and 25% (w/v) PEG1500, and mCherry-LaM6 complex crystallized in 100mM MIB buffer (Malonic acid, Imidazole, Boric acid (in a 2:3:3 molar ratio) pH 6.0 with 25% (w/v) PEG1500. The mCherry-LaM4 crystal was transferred into cryoprotectant that contains the crystallization condition supplemented with PEG1000 to 40% (w/v), while the cryoprotectant for mCherry-LaM6 crystal contains 100mM MMT buffer pH 6.0 with 25% (w/v) PEG1500 and 15% (w/v) glycerol. All crystals were flash frozen in liquid nitrogen prior to data collection. X-ray diffraction datasets were collected at the Advanced Photon Source on beamline 24-E. X-ray diffraction data were processed and scaled using the HKL2000 suite (25). Initial structures were solved using molecular replacement with a GFP (PDB: 4EUL) or mCherry (PDB: 2H5Q) structure as a search model using PHASER (26), and iterative rounds of model building in COOT (27) and refinement against the high-resolution datasets with PHENIX (28) were used to produce the final models. Diagrams of protein structure and RMSD calculations were performed using Pymol (Schrodinger).

### Thermal Shift Assay for nanobody stability

Nanobody stability was studied using a Protein Thermal Shift Assay (ThermoFisher). 0.5 mg/mL protein in 1X PBS was combined with 8X thermal shift dye and protein shift buffer following the manufacturers protocol. 1X PBS was used as a negative control and 0.5 mg/mL BSA was used to verify assay functionality. The protein mixtures and controls were placed in a real-time CFX96 PCR thermocycler (BioRad) and heated from 25° C to 99° C at a ramp rate of 0.3° C/ sec. The melting temperature was determined by taking the maximum of the melt curve derivative.

### Expression and purification of TOP2α using the mCherry tag

TOP2α (aa 1-1531) with an N-terminal TEV site was cloned into pDEST785, which encodes an N-terminal mCherry fusion using LR Clonase II reaction mix (Thermo) and TEV-TOP2α-pENTR, and expressed and purified essentially as described for YFP–tagged TOP2α (9). Briefly, 1.2 × 10^9^ HEK293F cells were transfected with 1.2 mg plasmid DNA and 1.4 mg PEI (Polyetheyleneimine, Polysci) in HyCell TransFx-H media (Cytiva) and grown for 72 hours. Cells were pelleted by centrifugation at 400 x g for 10 minutes, resuspended in 80 mL PBS, and pelleted again at 500 x g. Cells were lysed in 36 mL TOP2 lysis buffer (50 mM TRIS pH 8.0, 600 mM NaCl, 0.5% (v/v) NP-40 substitute (Sigma), and 1 mM TCEP) supplemented with Complete-EDTA free protease inhibitor cocktail (Roche), and sonicated with 3 cycles of 5 s sonication using a Branson sonicator set to 55% power, followed by 30 s cooling periods. Crude lysate was centrifuged at 25,000 x g for 10 minutes, then passed over a column with a 2.5 mL bed volume of pre-equilibrated LaM4 anti-mCherry sepharose resin. The resin was washed twice with lysis buffer, 4 times with wash buffer (50 mM TRIS pH 8.0, 400 mM NaCl, 0.1% (v/v) NP-40 substitute, and 1 mM TCEP), and twice with low salt wash buffer (20 mM TRIS pH 8.0, 300 mM NaCl, and 1 mM TCEP). Following these washes, the resin was washed quickly into TEV elution buffer (low salt wash with 20 μg mL-1 TEV protease), then stoppered and incubated overnight at 4 °C. TOP2 was eluted the next day by washing the resin with low-salt wash buffer. TOP2α was polished by sequential ion exchange chromatography (6mL Source 15S (Cytiva), 20 mM Tris pH7.5, 3mM DTT, and a 0–500 mM NaCl gradient) and size-exclusion chromatography (Superdex200 16/60 (Cytiva), 20mM Tris pH 7.5, 500 mM NaCl, 0.5 mM TCEP).

## Data availability

Atomic coordinates and structure factors have been deposited in the RCSB Protein Data Bank (PDB) under entries 7SAH (LaG16-eGFP complex), 7SAI (Lag30-eGFP complex), 7SAJ (LaM2-mCherry complex), 7SAK (LaM4-mCherry complex), and 7SAL (LaM6 complex).

## Author Contributions

A.T.Q.C.: methodology, investigation, validation, formal analysis, visualization, writing-original draft, writing-reviewing and editing. T.L.W.: investigation, analysis, visualization, writing-original draft, writing-reviewing and editing. M.J.S.: conceptualization, methodology, resources, supervision, writing-original draft, writing-reviewing and editing, funding acquisition.

## Supporting Information

This article contains supporting information with Tables S1–5 and Figure S1

## Acknowledgments

We would like to acknowledge M. Pillon (Baylor) and S. Andres (McMaster) for helpful comments and feedback about this project and manuscript. Funding for this project was provided by the Center for Biomedical Discovery and Mayo Clinic Startup funds to MJS.

## Declaration of interests

The authors declare that they have no known competing financial interests or personal relationships that could have appeared to influence the work reported in this paper.

## Notes

### Competing Interest Statement

The authors have declared no competing interest.

