## Supplementary data for "High Efficiency Recombinant Protein Purification Using mCherry and YFP Nanobody Affinity Matrices"

### Supplementary Materials

**Contains:**

Supplementary Tables 1-5
Supplementary Figure 1
…

**Supplementary Table 1.** Binding constants for the nanobodies used in this study as reported by the indicated reference

| Nanobody | $K_D$ (nM) | $K_{on}$ ($M^{-1}s^{-1}$) | $K_{off}$ ($s^{-1}$) | Reference |
|---|---|---|---|---|
| GFP-enhancer | 0.59 ± 0.11 | $2.45 \times 10^5$ (±0.2 %) | $1.45 \times 10^{-4}$ (±3.1%) | 9 |
| LaG16 | 0.69 | $1.6 \times 10^6$ | $1.1 \times 10^{-3}$ | 10 |
| LaG30 | 0.5 | $2.8 \times 10^6$ | $1.3 \times 10^{-3}$ | 10 |
| LaM2 | 0.49 | $3.6 \times 10^5$ | $1.8 \times 10^{-4}$ | 10 |
| LaM4 | 0.18 | $3.5 \times 10^5$ | $6.3 \times 10^{-5}$ | 10 |
| LaM6 | 0.26 | $4.8 \times 10^5$ | $1.2 \times 10^{-4}$ | 10 |

**Supplementary Table 2.** Comparison between nanobody yields from a 1 liter culture

| *Nanobody* | *Protein yield (mg)* |
|---|---|
| *GFP-enhancer* | 57.4 |
| *LaG16* | 5.2 |
| *LaG30* | 13.7 |
| *LaM2* | 3.2 |
| *LaM4* | 29.2 |
| *LaM6* | 3.5 |

**Supplementary Table 3.** Average melting temperatures for the nanobodies from thermal assays

| *Nanobody* | *$T_{melt}$ (℃)* | *±* |
|---|---|---|
| *GFP-enhancer* | 51.7 | 0.2 |
| *LaG16* | 46.8 | 0.2 |
| *LaG30* | 44.0 | 0.2 |
| *LaM2* | 56.5 | 0.2 |
| *LaM4* | 54.6 | 0.2 |
| *LaM6* | 44.8 | 0.2 |

**Supplementary Table 4. X-ray crystallography data collection and refinement statistics.**

| **Parameter:** | **Value:** | | | | |
|---|---|---|---|---|---|
| **Structure:** | **LaG16-eGFP** | **LaG30-eGFP** | **LaM2-mCherry** | **LaM4-mCherry** | **LaM6-mCherry** |
| **PDB Accession #** | **7SAH** | **7SAI** | **7SAJ** | **7SAK** | **7SAL** |
| Space group | $P222_1$ | I422 | C2 | C2 | $P2_1$ |
| Cell dimensions | | | | | |
| *a*, *b*, *c* (Å) | 75.3, 88.2, 132.3 | 173.2, 173.2, 92.3 | 171.4, 160.6, 88.8 | 156.5, 41.9, 47.8 | 74.1, 87.9, 74.6 |
| $\alpha, \beta, \gamma$ (°) | 90, 90, 90 | 90, 90, 90 | 90, 109, 90 | 90, 94, 90 | 90, 109, 90 |
| Wavelength (Å) | 0.97918 | 0.97918 | 0.97918 | 0.97918 | 0.97911 |
| Resolution (Å) | 44.12-1.60 (1.66-1.60) | 48.94-2.22 (2.31-2.22) | 71.00-2.37 (2.45-2.37) | 42.15-1.15 (1.19-1.15) | 49.97-1.93 (2.00-1.93) |
| $R_{merge}$ | 0.061(1.16) | 0.15(1.86) | 0.075(1.042) | 0.086(0.456) | 0.097(1.046) |
| $I / \sigma I$ | 31.3(1.5) | 18.7(1.7) | 18.2(1.4) | 21.8(4.3) | 19.3(1.7) |
| $CC_{1/2}$ | 0.999(0.544) | 0.998(0.504) | 0.997(0.385) | 0.997(0.910) | 0.990(0.528) |
| Completeness (%) | 99.60(96.37) | 99.16(95.40) | 99.09(99.14) | 96.95(76.69) | 98.42(95.06) |
| Redundancy | 6.8(5.7) | 13.6(13.9) | 3.6(3.6) | 9.6(5.9) | 7.4(5.7) |
| **Refinement** | | | | | |
| Resolution (Å) | 44.12-1.60 | 48.94-2.23 | 71.0-2.37 | 42.15-1.15 | 49.97-1.93 |
| No. reflections | 58084 | 34198 | 91214 | 106627 | 66659 |
| $R_{work}$ / $R_{free}$ | 0.1563/0.1768 | 0.1711/0.1974 | 0.2076/0.2341 | 0.1175/0.1445 | 0.1617/0.1921 |
| No. atoms (non-H) | 3356 | 3054 | 13653 | 3528 | 5948 |
| Protein | 2854 | 2785 | 13355 | 3007 | 5386 |
| Ligand/ion | 22 | 84 | 115 | 38 | 58 |
| Water | 480 | 184 | 183 | 483 | 504 |
| Average *B* factor (Å$^2$) | 32.58 | 65.50 | 71.96 | 16.03 | 43.60 |
| Ramachandran: Favored (%) | 99.14 | 98.27 | 97.12 | 98.21 | 98.80 |
| Allowed (%) | 0.86 | 1.73 | 2.82 | 1.79 | 1.20 |
| Outliers (%) | 0.00 | 0.00 | 0.06 | 0.00 | 0.00 |
| r.m.s. deviations | | | | | |
| Bond lengths (Å) | 0.007 | 0.009 | 0.005 | 0.013 | 0.005 |
| Bond angles (°) | 0.81 | 0.90 | 0.81 | 1.31 | 0.85 |

All data was collected from a single crystal.
*Values in parentheses are for highest-resolution shell (10% of reflections).

**Supplementary Table 5.** Binding interface areas between nanobodies and binding partners

| *Structure* | *Interface Area (Å²)* |
|---|---|
| *eGFP-GFP-enhancer* | 682 |
| *eGFP-LaG16* | 658 |
| *eGFP-LaG30* | 727 |
| *mCherry-LaM2* | 524 |
| *mCherry-LaM4* | 719 |
| *mCherry-LaM6* | 642 |

**Supplementary Figure 1.** (A) Melt curves for all nanobodies and (B) first derivatives

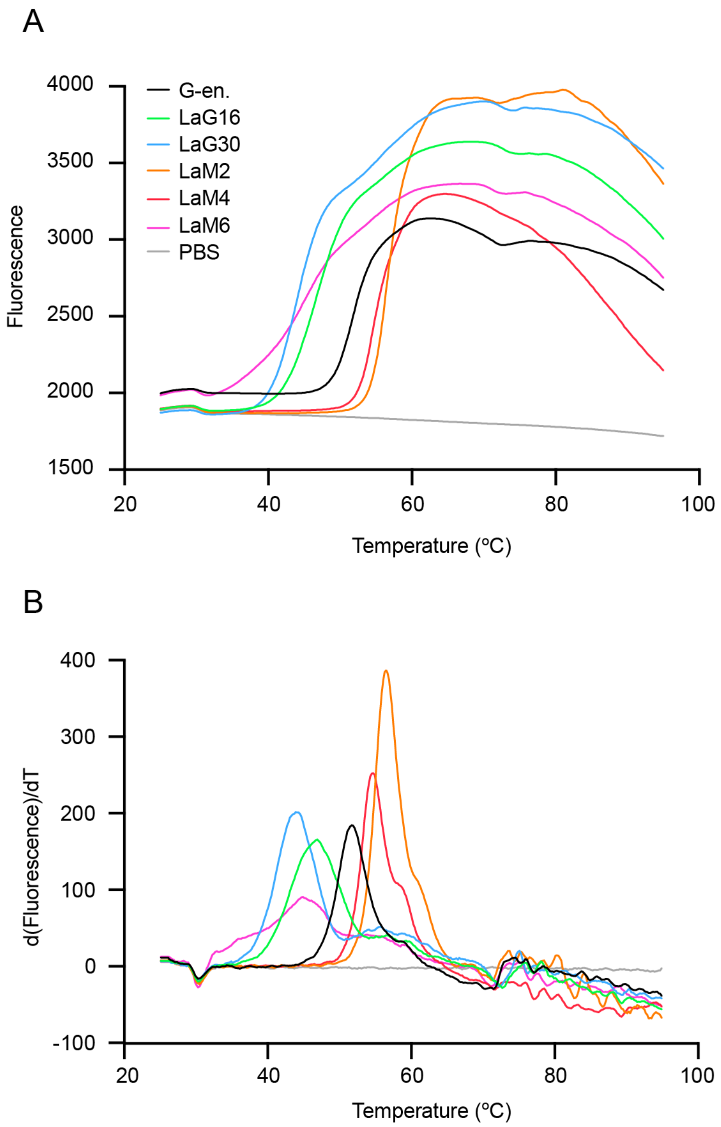

**Supplementary Figure 2.** Individual samples of fluorescent proteins that were used in resin binding assay. The anomalous band (*) is exclusive to the mCherry protein sample and present at a proportional fraction to the mCherry band, indicating it is a degradation product of mCherry.

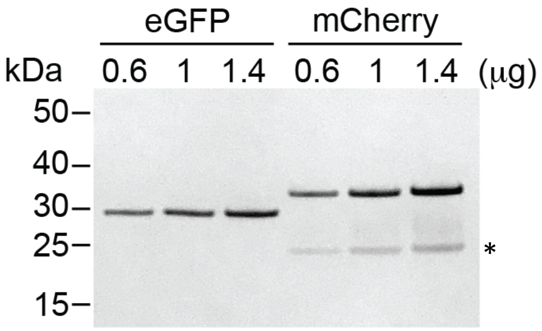
